# X-chromosome dosage compensation dynamics in human early embryos

**DOI:** 10.1101/2020.03.08.982694

**Authors:** Kevin Huang, Qiao Zeng, Yun Feng, Youjin Hu, Qin An, Taibo Li, Lian-Ju Qin, Jia-yin Liu, Zhigang Xue, Guoping Fan

**Author notes:** Corresponding authors: Guoping Fan, Ph.D., Department of Human Genetics, David Geffen School of Medicine, UCLA, 695 Charles Young Drive South, Los Angeles, CA 90095 USA, USA., And, Zhigang Xue, Ph.D., And, Jia-yin Liu, MD. These authors contributed equally to this project (Kevin Huang, Qiao Zeng, Yun Feng).

## Abstract

In mammals, female cells are obliged to inactivate one of two X chromosomes to achieve dosage parity with the single X chromosome in male cells, and it is also thought that the single active X chromosome is increased 2-fold to achieve dosage balance with two sets of autosomes (X:A ratio = 1, or Ohno’s hypothesis). However, the ontogeny of X-chromosome inactivation and augmentation of the single active X remains unclear during human embryogenesis. Here, we perform single-cell RNA-seq analysis to examine the timing of X:A balancing and X-inactivation (XCI) in pre- and peri-implantation human embryos up to fourteen days in culture. We find that X-chromosome gene expression in both male and female preimplantation embryos is approximately balanced with autosomes (X:A ratio = 1) after embryonic genome activation (EGA) and persists through fourteen days *in vitro*. Cross-species analysis of preimplantation embryo also show balanced X:A ratio within the first few days of development. By single-cell mRNA SNP profiling, we find XCI beginning in day 6-7 blastocyst embryos, but does not affect X:A dosage balance. XCI is most evident in trophoectoderm (TE) cells, but can also be observed in a small number of inner cell mass (ICM)-derived cells including primitive endoderm (PE) and epiblast (EPI) cells. Analysis between individual XaXa and XaXi sister cells from the same embryo reveals random XCI and persistently balanced X:A ratio, including sister cells transitioning between XaXa and XaXi states. We therefore conclude that the male X-chromosome undergoes X chromosome augmentation prior to the simultaneous X-chromosome inactivation and augmentation in females. Together, our data demonstrate an evolutionally conserved model of X chromosome dosage compensation in humans and other mammalian species.

## Introduction

Ohno’s hypothesis posits a two-step mechanism whereby X chromosome expression is increased two-fold in both sexes followed by X-inactivation in females(1). Various studies have tested Ohno’s prediction on a genome-wide scale, but they predominantly relied on somatic tissues and have led to conflicting results(2). We set out to re-visit the question of X chromosome dosage compensation in early human embryos by establishing cultures mimicking peri-implantation *in vitro*. Long-term culture of blastocyst embryos have been reported to resemble post-implantation development(3-5). Since we observed many blastocyst embryos hatching after 6-7 days in the standard medium in IVF clinic, we decided to co-culture hatched blastocyst embryos (6-days *in vitro*) with primary endometrial cells up to another eight days (**Fig. 1a**). This allowed us to obtain viable embryonic cells between 6 and 14 days *in vitro*.

**Fig. 1.**
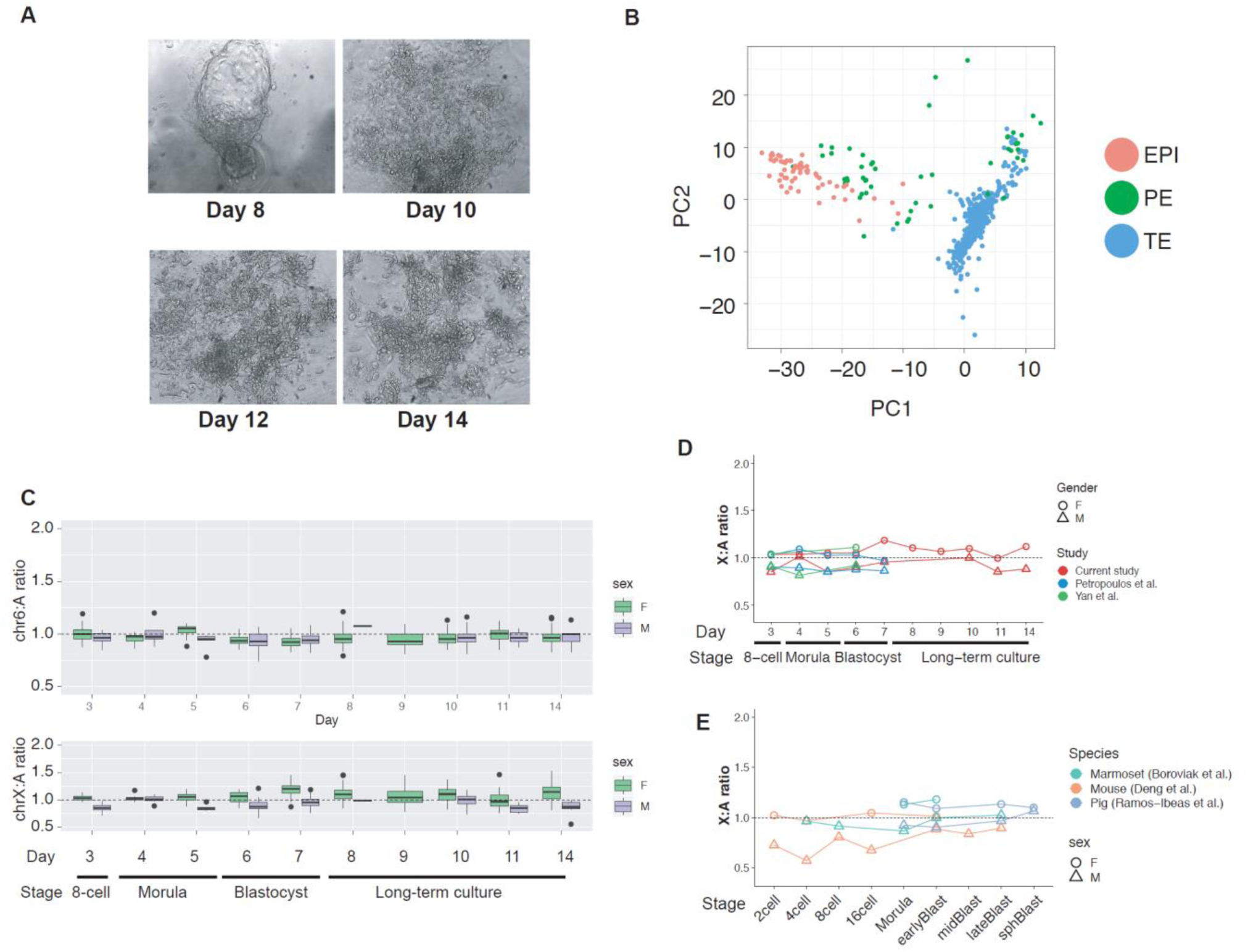
X:A dynamics in mammalian development. **a)** Brightfield image of human embryos in long-term culture and co-cultured with primary endometrial cells.**b)** Clustering of late stage embryos in their expression of 245 marker genes from day 6 to day 14 *in vitro*. Each dot represents a single-cell and the color is based on their lineage classification as determined by their expression of lineage-specific marker genes (see **materials and methods**). C) Boxplot showing the distribution X-to-autosome (X:A) ratios in single-cells across different days of in vitro culture. D; E) Meta-analysis of X:A ratios across multiple studies. Each point represents the median X:A ratio in each study at a particular time point (x-axis) for either female (open circle) and male (open triangle).

## Results

To gain a high resolution study of the X chromosome dosage compensation process in human embryos, we performed single-cell RNA-sequencing in 73 embryos for a total of 953 single-cells **(Supplementary Table S1)**. To remove potential bias result from aneuploidy, we excluded 248 cells with at least one aneuploid chromosome arm, as well as 77 syncytiotrophoblasts. In addition, we re-analyzed over 1,600 single-cell RNA-seq in human pre-implantation embryos from previous studies(6, 7). Leveraging previous marker genes for EPI, PE, and TE lineages(6, 8), we were able to discriminate all three lineages in long-term cultured human embryos (**Fig. 1b and Supplementary Fig S1**).

We carried an analysis that compared the median expression ratio between the X chromosomes and autosomes (X:A) as a measure of X-to-autosome dosage compensation(9-12). In our analysis, both female and male preimplantation embryos exhibit a nearly balanced X:A ratio (0.93±0.01 in males and 1.09±0.01 in females, mean±s.e.m.) throughout all stages of long-term culture beginning from the 8-cell stage (**Fig. 1c)**. These data suggest that the transcriptional output from the X chromosome is on par with autosomes upon embryonic genome activation (EGA). A corollary of our finding is that females and males have balanced X chromosome expression, indicating that the single male X-chromosome exhibits approximately 2-fold greater transcriptional output compared to each active female X chromosome after EGA. Control comparisons with other autosomal chromosomes (chr1, chr2, chr6, chr7) also reveal balanced dosage (**Supplementary Fig S2**). There were also no differences in X:A ratios between the three lineages (**Supplementary Fig S3)**.

We re-analyzed a previous single-cell RNA-seq dataset of human preimplantation and confirmed approximately balanced X:A dosage balance in both males and females embryos (0.88±0.01 in males and 1.03±0.01 in females, **Fig 1d and Supplementary Fig S4a**). Consistent with a previous re-analysis(13), our findings are in contrast to the proposed model of X-chromosome “dampening”, whereby female cells in cleavage embryos transiently express cumulatively 2-fold higher transcription of the X-chromosome compared to males and gradually reach parity in blastocyst embryos. Our approach to take the median of expressed X-linked genes is likely more resistant to over-amplified outlier genes compared with the previous method to sum all expressed genes on the X-chromosome. We re-analyzed a third dataset(7) and found similar results (**Fig 1d and Supplementary Fig S4b**). Together, these results indicate that the X-to-autosome ratio is rapidly established in both sexes upon EGA and persists through all stages of pre- and peri-implantation.

To test for evolutionary conversation of the X:A dosage compensation, we analyzed single-cell RNA-seq of preimplantation embryos from multiple species, including non-human primates(14), pig(15), and mouse(16). In general, we observed similarly balanced X:A dosage early in development. For example, marmoset males exhibit X:A ratio of 0.97±0.05 during the 4-cell stage, and maintains a near equivalent X:A ratio for the rest of preimplantation development (**Fig. 1e**). In pig, males and females show similar X:A ratio at the morula and blastocyst stages (0.97±0.02 in males vs 1.12±0.03 in females). In mice embryos, we observed that male embryos showed lower X:A ratio in cleavage embryos (ranging between 0.57 and 0.81) but becomes nearly balanced in blastocyst stages (0.87±0.03). However, female embryos shows consistent balanced X:A ratio through all stages (1.01±0.05) (**Fig. 1e**). Together, these data suggest that X:A dosage compensation is achieved by morula and blastocyst stages of development, suggesting a conserved mechanism of X-chromosome dosage compensation in evolution.

We next investigated X-inactivation (XCI) in our culture system to model per-implantation human embryonic development. XCI is another mode of X-chromosome dosage compensation that provide X-chromosome parity between the sexes(1, 2, 9, 10, 17, 18). Previous studies have suggested that X-inactivation in human embryos is distinct from that found in the mouse model system(19-21). For example, human *XIST* expression is first detected in the 8-cell stage(22, 23) and coats both X-chromosomes in morula and blastocyst cells without apparently triggering XCI(19). The lack of XCI in human preimplantation embryos was recently confirmed via single cell RNA sequencing analysis^5^ but refuted by a recent re-analysis of the same dataset(13).

We analyzed global X-chromosome silencing in our culture system by leveraging mRNA SNPs to estimate the number of bi-allelically expressed X-linked genes in female cells across the entire X chromosome (see Methods). We reason that cells that have undergone chromosome-wide silencing would exhibit significantly fewer biallelically expressed X-linked transcripts compared to autosomes (**Fig. 2a, one-way binomial-test adjusted-p<0.05**). Using male embryos and endometrial cells as reference, our single-cell analysis SNP analysis identified a portion of single-cells from female embryos that resemble monoallelic X chromosome expression, indicating that XCI onset can be modeled in human peri-implantation embryos (**Fig. 2b**). We also note that the average proportion of biallelically expressed X-linked SNPs show a continuous distribution, suggesting that XCI is a gradual process (**Fig. 2b**). Because of this spectrum, we separated female single-cells into three categories: bi-allelically expressing X (XaXa), partially inactivated (pXCI), or mono-allelically expressing X (XaXi). Remarkably, the X-to-autosome ratio between the three states is fairly balanced. In aggregate, the X:A ratio is 1.14±0.13 (mean±s.d.) in XaXa cells, 1.11±0.13 in pXCI and 1.07±0.10 in XaXi cells (**Fig. 2c**), which suggest that the two-fold up-regulation of X chromosome likely occurs rapidly during the XCI process.

**Fig. 2.**
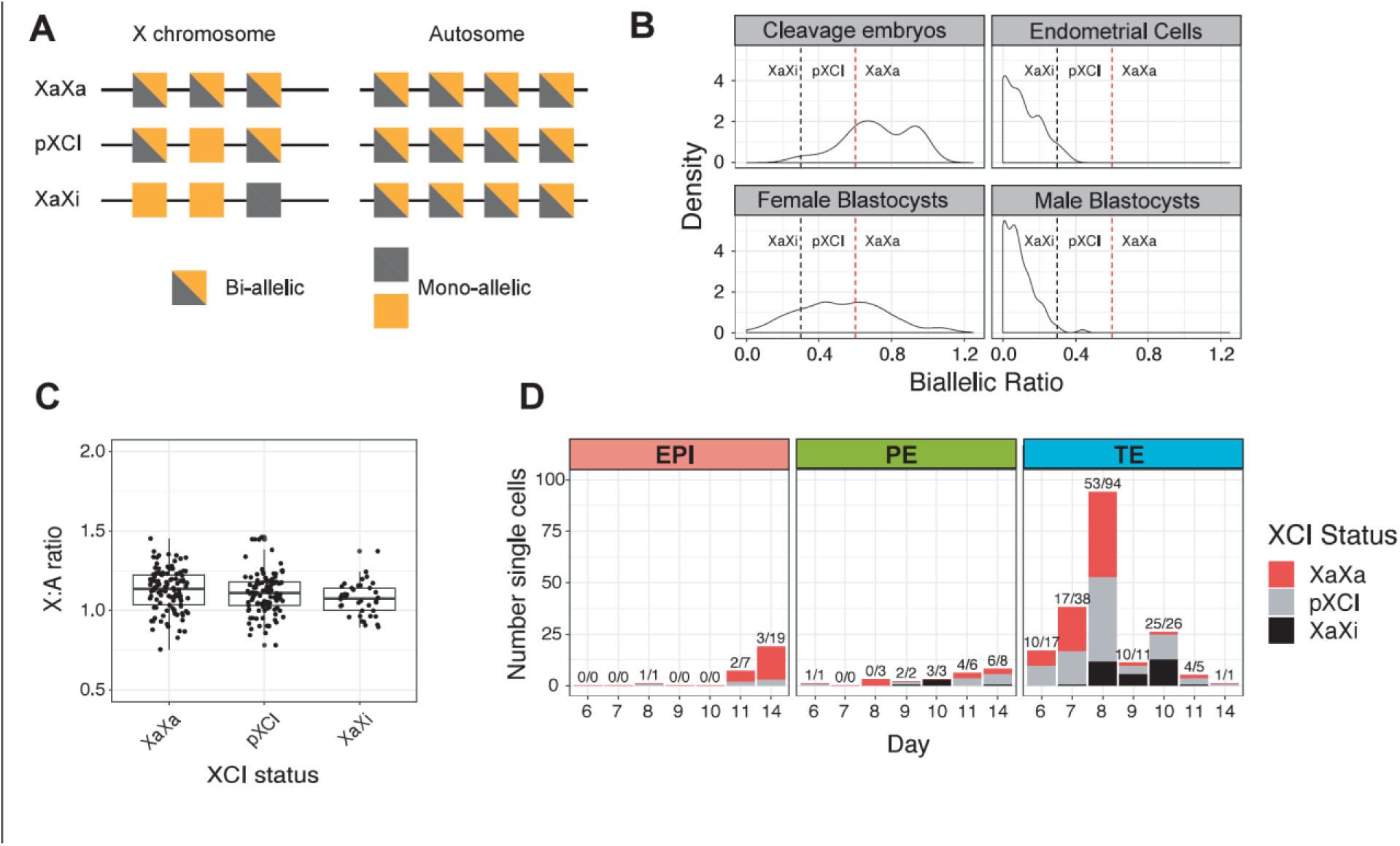
X-inactivation dynamics in human pre- and peri-implantation embryos. a) Schematic showing different scenarios for determining XaXa vs XaXi classification. b) Density plot showing distribution of biallelic ratio, (the fraction of X-linked biallelic SNVs vs fraction biallelic autosome SNVs). c) Boxplot showing the distribution of X:A ratio from female embryonic cells carrying different X-inactivation status (x-axis). D) Stacked barplot showing the distribution of X-inactivation states across lineages and across developmental days. Number above the bars represent the number of partial or complete XCI cells and the total number of cells.

Across different cell lineages, XCI is most pronounced in TE cells but not in EPI nor PE cells (**Fig. 2d**). While this finding is consistent with previous report of greater number of TE cells undergoing XCI in long-term human embryo culture(24), our data is also limited due to under-sampling of EPI and PE cells. For example, we were able to collect 30 EPI cells, of which 6 showed partial X-inactivation and 0 showed full XCI. By contrast, we collected 201 female TE cells of which 92 (46%) were pXCI and 33 (16%) were complete XCI. TE cells showed progressively greater number of cells with XCI over time with the greatest change after day 7 where the average proportion of XCI cells changed from 5% to approximately 50% in day 9 and day 10 embryos (**Fig 2d**). These data suggest that TE cells preferentially undergo XCI.

In our re-analysis of a larger set of human preimplantation data(6), we also found similar XCI findings. Firstly, we observed a common continuous distribution of the biallelic ratio between X chromosome and autosomes. A small proportion of female cells were classified as pXCI and XaXi (**Supplementary Fig. S5a**), consistent with a previous re-analysis(13). These XCI cells were either EPI or TE (**Supplementary Fig. S5b**). Thus, XCI states can be identified as early as day 6. Overall, these results indicate that XCI first takes place in 6-day old embryos, particularly in TE, with gradually increasing number of cells in the following week in culture.

We next searched for sister cells from the same embryo which contained all three states of X-inactivation (XaXa, pXCI and XaXi). This offered us the powerful ability to examine the dynamics of X chromosome dosage compensation of genetically identical and developmentally synchronized cells. We identified two female day 8 embryos from which a sufficient number of XaXa, pXCI, and XaXi sister cells were represented. However, this analysis was restricted to TE cells due to insufficient number of EPI and PE cells from the same female embryo. Strikingly, XaXa, pXCI, and XaXi sister cells all showed balanced X-to-autosome expression (**Fig. 3a**). Analysis of individual genes between XaXa and XaXi states also revealed that the vast majority of genes were not significantly differentially expressed, suggesting that XCI is quickly compensated with 2-fold output from the single active X chromosome (**Fig. 3b and Supplementary Fig. S6**). While no genes were statistically significantly differentially expressed, we found that only *XIST* expression in XaXi cells were consistently upregulated compared to their XaXa sister cells in both embryos (nominal p-value < 0.03, **Fig. 3c**.), in line with the classic model of allelic *XIST* upregulation triggering XCI. *TSIX* expression was not robustly expressed enough to perform meaningful analysis.

**Fig. 3.**
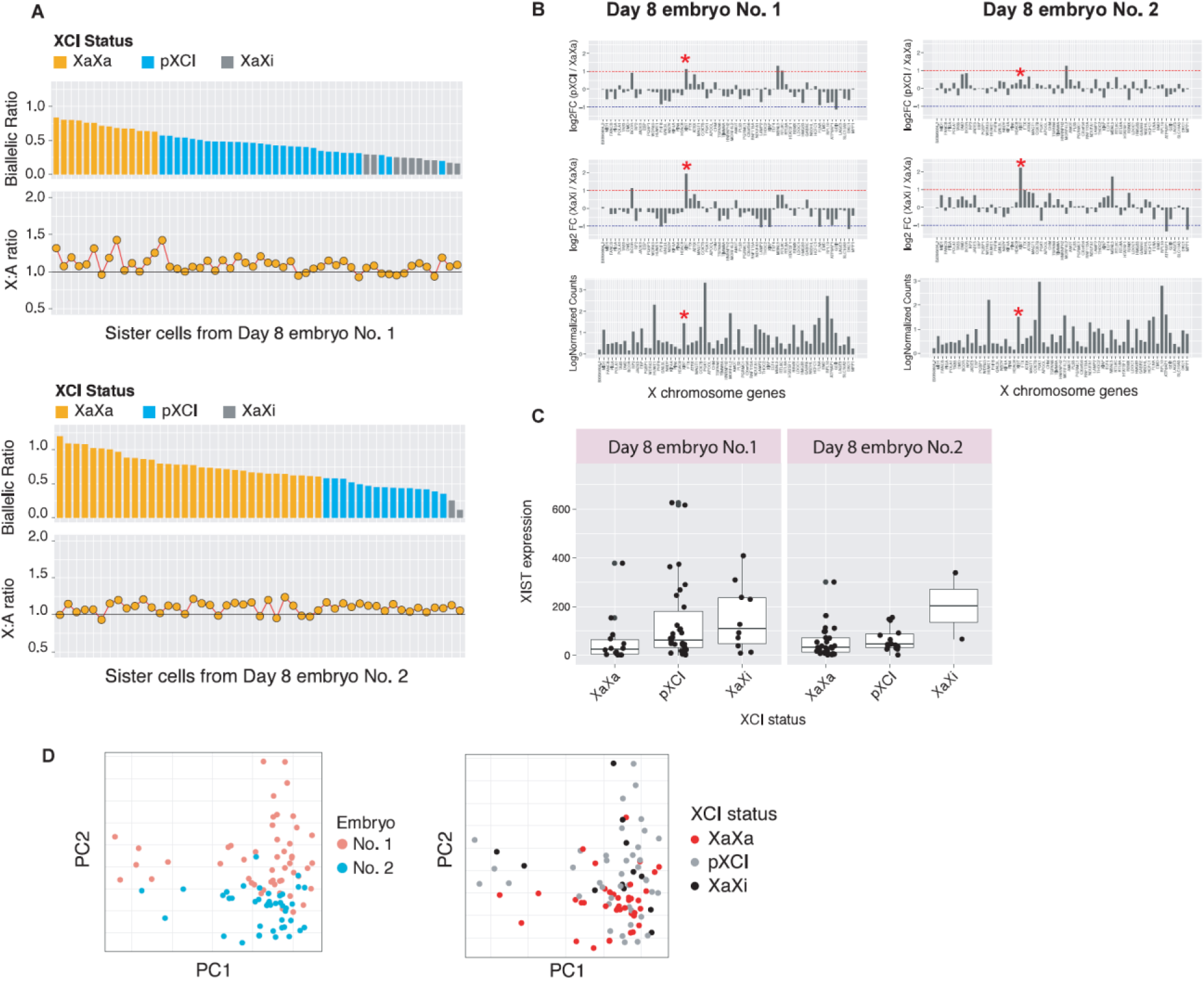
Sister cell analysis X chromosome dynamics. a) Top: Barplot of sister cells from the same embryo ranked by their biallelic ratio. Bottom: Corresponding X:A ratio in each single-cell. B) Barplot showing the median log2 fold change of genes between pXCI vs XaXa (b) or XaXi vs XaXa sister cells. Bottom panel shows the average gene expression. E) heatmap showing the percent of cells that show biallelic expression of a gene. F) Boxplot of XIST expression from sister cells with different X-inactivation states.

It is thought that XCI is associated with many accessory transcriptome-wide changes, such as downregulation of pluripotency factors and upregulation of differentiation genes(25). Clustering analysis revealed no clear separation between XaXa, pXCI, and XaXi sister cells, consistent with our differential gene expression analysis (**Fig. 4a**). This suggests that XCI occurs independently of any major genome-wide transcriptional changes.

**Fig. 4.**
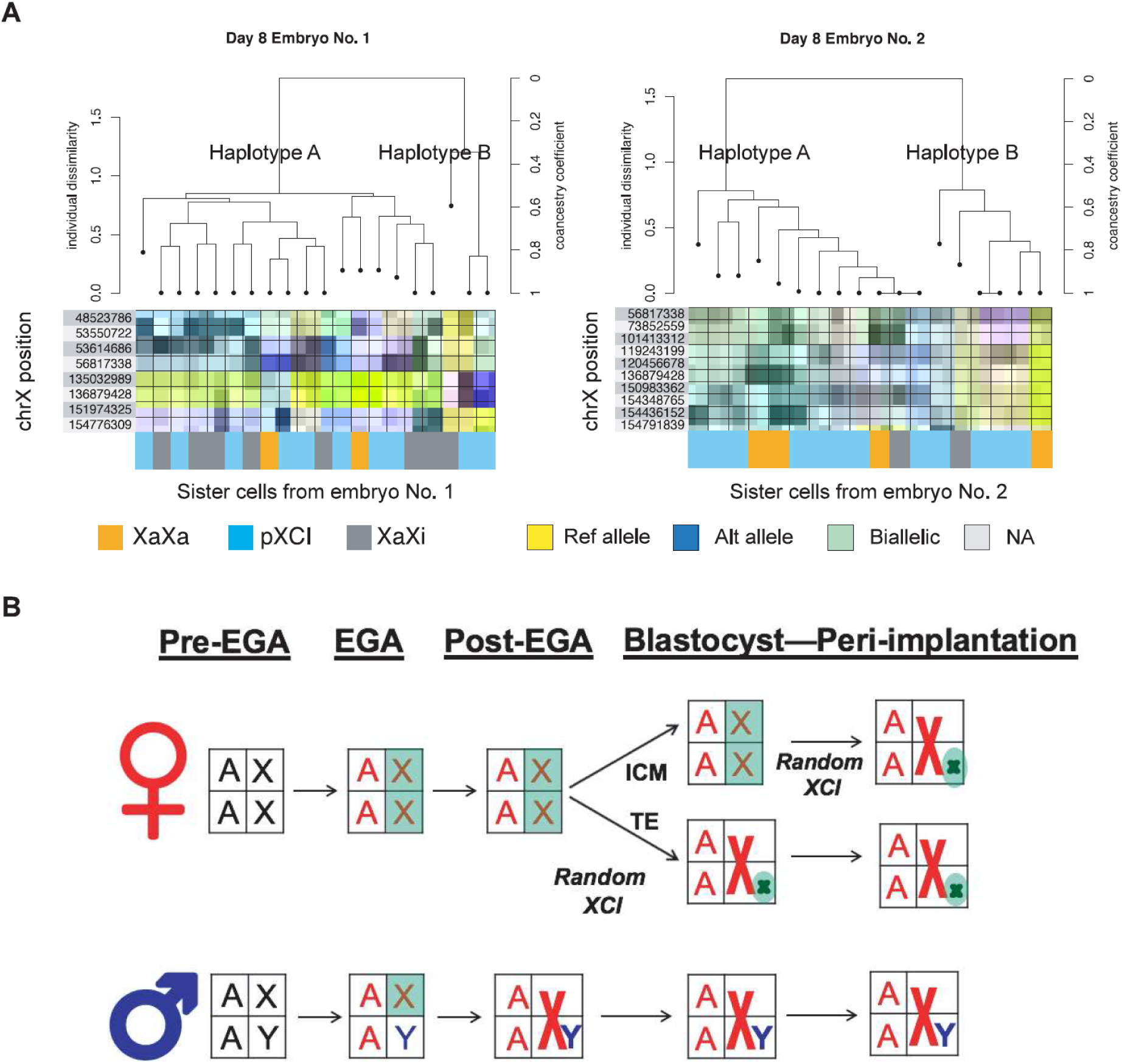
Model of X chromosome dosage compensation in early human development. a) PCA plots of sister embryos colored by their X-inactivation states b) PCA plot of sister embryos based on their inferred genotypes from single-cell RNA-seq data c) barplot showing distribution of haplotypes in each individual embryo and X-inactivation state (x-axis) d) Schematic model of X-chromosome dynamics in pre-implantation and post-implantation embryos.

Finally, we were able to phase a small number of SNPs along the X chromosome between sister cells to determine whether X chromosome silencing occurred randomly (see Methods). In both day 8 embryos, we were able to identify two haplotypes (**Fig. 4b**). One embryo had sufficiently more XCI skew (81% of haplotype A and 19% of haplotype B) whereas another embryo had more even number of random XCI (67% haplotype A and 33% haplotype B) (**Fig. 4c**). Together, these data suggest random XCI.

## Discussion

Our single-cell analysis of the X-chromosome transcriptome in long-term co-cultures of human embryos with endometrial cells uncovered new insights into X-chromosome dosage compensation *in vitro* **(Fig 4d)**. X chromosome parity between the sexes is achieved through a combination of XCI and augmentation of active X-linked gene expression. In male embryonic cells, the single X chromosome gene expression is augmented by approximately two-fold immediately after the first wave of EGA. In female cells, EGA triggers two active Xs that are balanced with autosomes, but random XCI can take place from blastocyst embryos to peri-implantation period in vitro. While the timing of random XCI in different female cells can be variable over eight days *in vitro*, we found that dosage compensation of X:A is balanced at 1:1 regarding of preXCI, partial XCI or post-XCI. The molecular mechanism involved in sensing X:A dosage compensation is still unclear, but it is likely triggered by evolutionarily conserved regulatory feedbacks due to the ratio of active X:A gene expression. While XCI dynamics vary across eutherians(19), the mechanisms of X:A dosage balancing is likely evolutionarily conserved. In the derivation or long-term cultures of human embryonic stem cells (hESCs) or induced pluripotent stem cells (hiPSCs), we and others also found that hESCs/hiPSCs can exhibit complex XCI regulation including random or skewed XCI, the erosion of XCI upon long-term cultures or reversible reactivation and silencing of the inactive X under naïve or prime pluripotent state(26, 27). These findings are consistent with the notion that human embryonic cells may adapt to different states of XCI pending under specific physiological or pluripotent states in vitro. Future strategies such as CRISPR/Cas9 mediated screening of X-linked or autosome genes in hESCs/hiPSCs or in non-human primate models may aid to understand the fundamental mechanism of X dosage compensation mechanisms during early primate embryogenesis.

## Materials and Methods

### Ethics Statement

Human preimplantation embryos (2-cell to blastocyst embryos) were obtained from female patients (between ages 26 to 35) at the Center for Clinical Reproductive Medicine at the Jiangsu People’s Hospital/First Affiliated Hospital of Nanjing Medical University through written informed consent and institutional approval. This study was approved by the Institutional Review Board (IRB) on Human Subject Research and Ethics Committee in the First Affiliated Hospital to Nanjing Medical University in China. None of the donors received any financial re-imbursement. Patient IDs were anonymous to all the scientists in this project. Investigators at UCLA were only involved in data analysis and manuscript writing. According to UCLA IRB review of their involvement in the study, UCLA investigators were granted exemption approval (IRB#12-001361).

### Human embryo collection and culture

Human embryos collection was performed as described previously1. Briefly, embryos were cultured in Cleavage Medium (SAGE, USA) in a low-oxygen humidified atmosphere containing 5% CO2, 5% O2, and 90% N2 at 37°. Stages of human embryo development were assessed through microscopy and collected separately.

For the peri-implantation culture, vitrified day3-embryos were warmed with a thawing kit according to the manufacturer’s instructions (Jieying Laboratory). All thawing steps were performed at room temperature (23–25 °C). Embryos were thawed by placing the cryovials in an in TS1 for less than 3 min at room temperature, followed by incubation in TS2 and TS3 sequentially for 3 minutes at room temperature. Embryos were cultured in blastocyst Medium (SAGE) in a low-oxygen humidified atmosphere containing 6% CO2, 5% O2 and 89% N2 at 37 °C and stages of embryo development were assessed through microscopy and collected separately. Hatched blastocysts were morphologically scored after culture for 72 hours. Zona was removed by exposing to acidic tyrode’s solution (Sigma). Zona-free blastocyts were seeded on endometrial cells and cultured in DMEF/F12 medium with 10% human serum albumin in a low-oxygen humidified atmosphere containing 6% CO2, 5% O2 and 89% N2 at 37 °C. Zona-free blastocysts typically attached 48–60 h after seeding. Half of the medium was replaced with fresh medium every 24 hours.

### Single cell isolation

To collect single cells from non-attached blastocysts, embryos were treated with Tyrode’s acidic solution to remove the zona pellucida then seperated by 0.25% typsin digestion. Single blastomeres were washed twice with 1xPBS containing 0.1% BSA and transferred to Smart-seq2 lysis buffer (0.2% Triton-X100, 1U RNase inhibitor, 2.5 μM oligo dT primer and 2.5 mM dNTP) by mouth pipetting.

To collect single cells from attached embryos, the attached embryos were first separated from the endometrial cells by gentle mouth pipetting, and then transferred to drops of 0.25 % trypsin-EDTA. Single cells were separated by repeated pipetting and washed twice with 1×PBS containing 0.1% Polyvinylpyrrolidone before placing in Smart2-seq lysis buffer.

### RNA isolation and library construction

Whole-transcript amplification for single blastomeres was done using the SMART2 protocol2. Briefly, cDNA was first generated by reverse transcription with SuperScript III (Life technologies, 18080-044). Binding sequences of primers for cDNA amplification which were anchored at the oligo dT primer and LNA-TSO primer were added to both end of cDNA during reverse transcription. Amplification of cDNA was performed by using the KAPA HiFi HotStart ReadyMix (KK2602) and the PCR products were purified by using Ampure XP beads (1:1 ratio). Libraries were constructed by using llumina Nextera XT DNA sample preparation kit (FC-131-1024) with 0.1ng cDNA. Library quality was assessed using Agilent Bioanalyzer 2100 and sequencing was performed in either a Illumina Hiseq 2000 or Illumina Nextseq 550 machine.

### Read alignment and QC

Raw reads were first trimmed using Trim Galore with paramaters --q20 --phred 33 --length 30. Raw reads were mapped to the human (hg38) genome and using default parameters in the STAR aligner3 (version 2.7.3g) using default parameters. The primary genome and gene annotation models were downloaded from Gencode (version 29).

To reduce mapping error due to polymorphic sites, we used the WASP software4, which is a pseudo-SNP aware method for re-alignment, to remove reads that could be erroneously mapped due to single nucleotide variants. Only reads that mapped to the same mapping position after taking into consideration common SNPs were retained. SNPs.

Our panel of SNPs were downloaded from the 1000 genomes database (March 2019 release, http://ftp.1000genomes.ebi.ac.uk/vol1/ftp/data_collections/1000_genomes_project/release/20190312_biallelic_SNV_and_INDEL/). For our study, we restricted our analysis to common SNPs and indels (minor allele frequency > 1%) from the East Asia population. For the Petropoulos et al dataset re-analysis, we used the identical workflow except we included common European SNPs (MAF > 1%). In all analyses, only reads that were uniquely aligned were retained.

### Sample processing

A raw count matrix were generated using the featureCount5 function in the subread package (v1.6.4). For subsequent analysis including data normalization, visualization, lineage classification and gender determination we used the R software package Seurat (version 2.3.4). Specifically, read count matrix was converted into a Seurat object using default parameters. The Seurat object were then normalized using Seurat NormalizeData function (parameter scale.factor was set to 10000) and scaled using ScaleData function, to regress out the sequencing depth variation.

Single cells with mitochondrial reads that exceeded 25% of total reads were excluded. Using these criteria, 790 out of 891 (88.7%) single cells from 42 embryos (Supplementary Table S1). Differential expression analysis was performed using the Seurat function FindMarkers with default Wilcoxon Rank Sum test.

### Lineage classification

Lineage classification were performed using customized R script modified from Seurat CellCycleScoring function, which is available online (https://github.com/Winbuntu/CellLineageScoring/blob/master/function.R). The principle of this method is described in another study6. Briefly, we first collected 300 maintained marker for EPI, PE and TE lineages. For each cell, we computed a “TE score”, an “EPI score” and a “PE score” were computed using AddModuleScore function implemented in Seurat package, based on its expression of markers for each lineage, respectively. The linage of a cell was then defined as the lineage that yield the highest score.

### Embryo sex determination

The gender of each single cell was determined by their expression of Y chromosome-linked genes. Briefly, the sum counts of reads mapped to all chrY linked genes for each cell were normalized by total read depth. The normalized chrY read counts for all cells were then log-transformed. The normalized, log-transformed chrY read counts exhibit bimodal distribution, representing the male and female cells respectively. The gender of each cell was determined based on their normalized, log-transformed chrY read counts (Supplementary Fig S1 and Supplementary Table S1). After determining the gender of each cell, the gender of each embryo was determined according to the consensus gender of all cells from their respective embryo. Embryos containing less than 3 cells were or contained significant number of both male and female cells were excluded from X chromosome analysis. We also further validated gender by examining XIST expression which is predominantly expressed in female cells. For a subset of male cells, we did detect substantial levels of XIST, which have been observed before7, but it remains unclear how to interpret these samples. These samples with ambiguous gender were excluded from our X-chromosome analyses.

### Variant calling

SNP processing and calling was performed using Picard (v2.18.29) and GATK (v3.8-1-0-gf15c1c3ef)8. Each cell was processed individually. Reads marked as PCR duplicates were removed with Picard’s MarkDuplicates function and then re-assigned mapping qualities using the Split’N’Trim function and BaseRecalibrator function using default parameters.

Variant calling was performed using the HaplotypeCaller function with parameters -dontUseSoftClippedBases -stand_call_conf 20.0. Only SNPs with QD score > 20 and covered by at least 8 reads or more were retained for further downstream analysis. Allele counts were retrieved using the ASEReadCounter (v3.7) function for SNPs that passed VQSR filtering using parameters --minMappingQuality 60 -- minBaseQuality 10 -drf DuplicateRead -minDepth 8.

### SNP background correction and filtering

We noted that there were spurious biallelic sites in male samples were likely due to noise which could confound our estimation of biallelic sites on the X chromomosome. To adjust for noise at each SNP, k, we first derived a population average of allelic balance (AB, defined as the number of max(reference counts, alternative counts)/(ref + alt counts), p, from all male samples for each position. For each cell at position k, we performed a one-sided binomial test Binom∼(x,n,p) where x is the larger of reference or alternative counts, n is the total (ref+alt) counts, and p is the male population average AB. SNPs which had a nominal p-value < 0.05 were considered biallelic, else regarded as monoallelic expressed.

SNPs that were within known escape genes were excluded from analysis. We used escape genes described by Tukiainen and colleagues9 which surveyed escape genes across hundreds of tissue samples in the GTEx database as well as in single-cells. In total 96 escape genes were excluded.

### XCI status inference

Each cell was required to carry at least 40 X-linked SNPs to be included the analysis, though we arrive at similar conclusions when raising or lowering this arbitrary threshold. For each individual cell, we used a one-sided binomial test Binom∼(x,n,p) where p is the proportion of heterozygous SNPs across all autosomes, x is the number of heterozygous SNVs on the X chromosome, and n is the total number of X-linked SNPs. This produced a p-value which informed the rejection of the null hypothesis that the X chromosome follows a biallelic expression analogous to autosomes. P-values were adjusted for multiplicity using the Holm method, and cells which had adjusted p-values < 0.05 were considered to monoallellic expressing the X chromosome(s). Nearly all male samples were called monoallelic and the vast majority had a bi-allelic ratio of less than 0.3. Therefore, we also applied this threshold of 0.3 for female samples in addition to the adjusted p-value threshold to further increase stringency. Female cells were considered bi-allelically expressing X chromosomes if their bi-allelic ratio was greater than 0.6 and nominal p-value > 0.05. The threshold of 0.6 was chosen based on comparison with cleavage embryos which are known to have two active X chromosomes. Cells that did not satisfy either XaXa or XaXi criteria were considered partially inactivated (pXCI) (i.e. most cells that fell within the 0.3 and 0.6 range).

### Single-Cell RNA Seq Aneuploidy Calling

To infer copy number changes on the chromosomal scale from single-cell transcriptomics data, we leveraged the fact that quantitative change in chromosome content can be reflected as gene expression dosage changes calculated on each chromosome. We first applied deep count autoencoder (DCA) to denoise single-cell RNA-seq count matrix, which has demonstrated reliable performance in dissecting cell population structures in real and simulated datasets10. We then studied EPI, PE, and TE cell populations separately, and calculated a chromosome score for each chromosome in each cell characterizing how much normalized total gene expression on that chromosome deviates from the median of their respective populations, as in Griffiths et al11. We subsequently called aneuploidy based on predefined threshold (FDR corrected q < 0.1) similar to Griffiths et al11.

### Estimation of X-to-autosome expression ratios

X chromosome expression was calculated by taking median value of all highly expressed X-linked genes (logNormalized counts > 0.01). Pseudoautosomal and X-linked escape genes as defined by Tukiainen et al9. were excluded from analysis. On average, we consistently detect between 200-300 X-linked genes per cell (Supplementary Table S1). Median autosome expression was calculated by taking median value of all highly expressed autosomal genes (logNormalized counts > 0.01). The X-to-autosome (X:A) ratio was calculated by dividing the median X chromosome expression with median autosome expression.

### Random XCI inference

For each embryo, we first filtered for SNPs that has monoallelic reference allele and monoallelic alternative allele represented by at least one cell. We next required that for each individual cell to carry at least 2 monoallelically expressing SNPs. This procedure greatly reduced our search space for relevant SNPs for phasing. We generated a genotype matrix of n SNPs x p samples. which was used the SNPRrelate R package12 for LD pruning, hierarchical clustering, and estimating kinship coefficients. A z-score to estimate significance was performed by 10,000 rounds of permutation testing.

### Data visualization

Data visualization such as PCA was performed using Seurat built-in functions and customized R scripts. Heatmaps were visualized using the superheat package (https://rlbarter.github.io/superheat/). Haplotype dendrograms were visualized using the SNPRelate R package12.

## Acknowledgements

We thank many of our colleagues for invaluable discussions and comments on our study. The work was supported by National Program on Key Basic Research Project (973 Program); National Natural Science Foundation of China (Key Program 81430026, Youth Program 31301184); Jiangsu Province Clinical Research Center (BL2012009).

## Author contributions

KH, QZ, ZX, and GF designed the study. KH and QA analyzed the data and performed statistical analyses. TL performed the aneuploidy analysis. FY, QZ, YH, TL, LQ, ZX, JL performed experiments or contributed critical reagents and protocols. KH and GF wrote the manuscript in discussion with all authors. All authors read and approved the manuscript.

## Author information

All sequencing data generated for this work have been deposited in the NCBI Gene Expression Omnibus (GEO GSE 130050) and would be released upon publication of this paper. The authors declare no competing financial interests. Correspondence and request for materials should be addressed to GF, and ZX and JL.

## Supplementary Figures

**Supplementary Figure S1.**
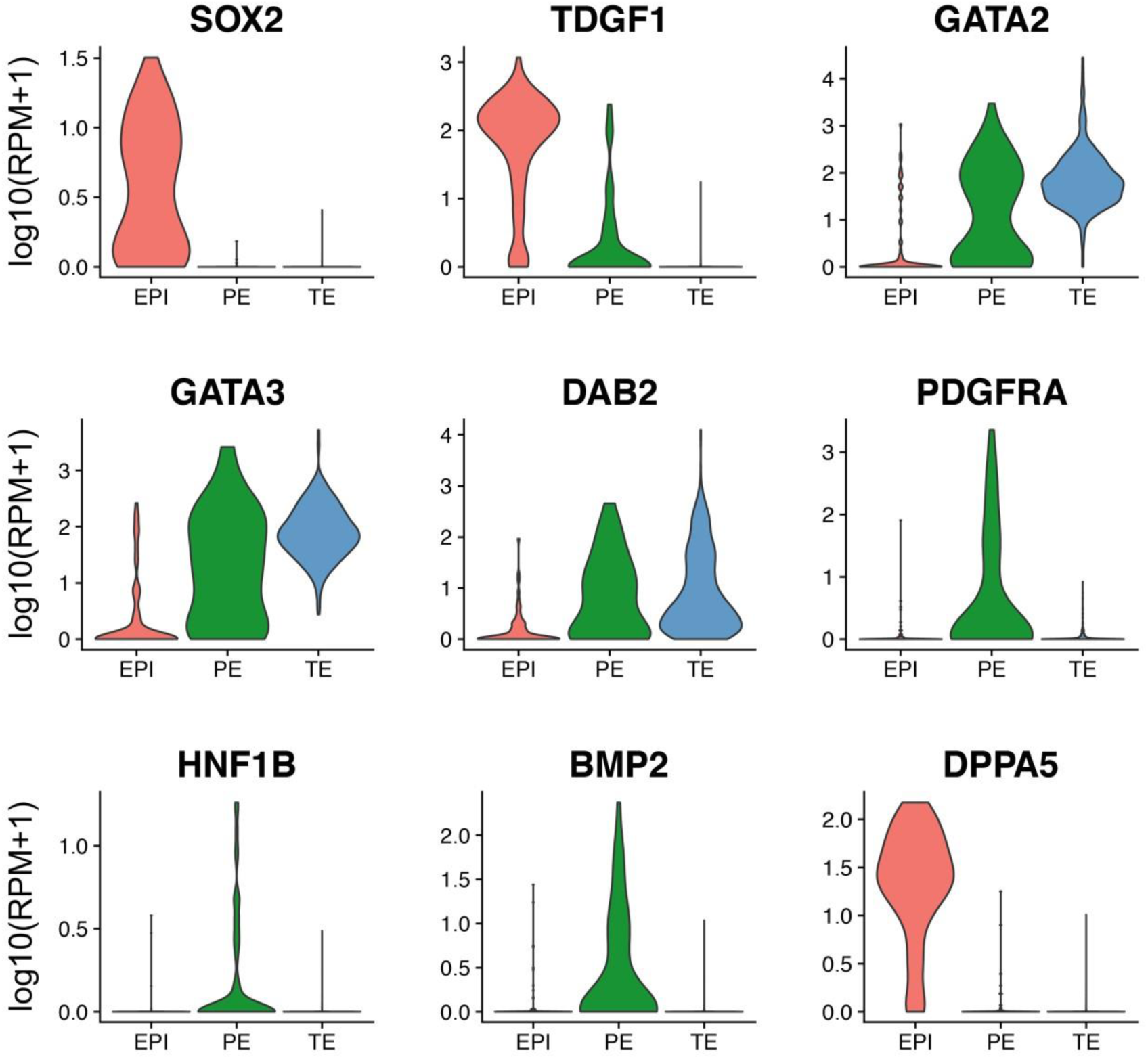
Lineages of single-cells from embryos can be discriminated. Violin plots of example marker genes used in our inference of cell lineage (x-axis). EPI: epiblast; PE: primitive endoderm; TE: trophoblasts. Y-axis shows the lognormalized expression values from Seurat.

**Supplementary Fig S2.**
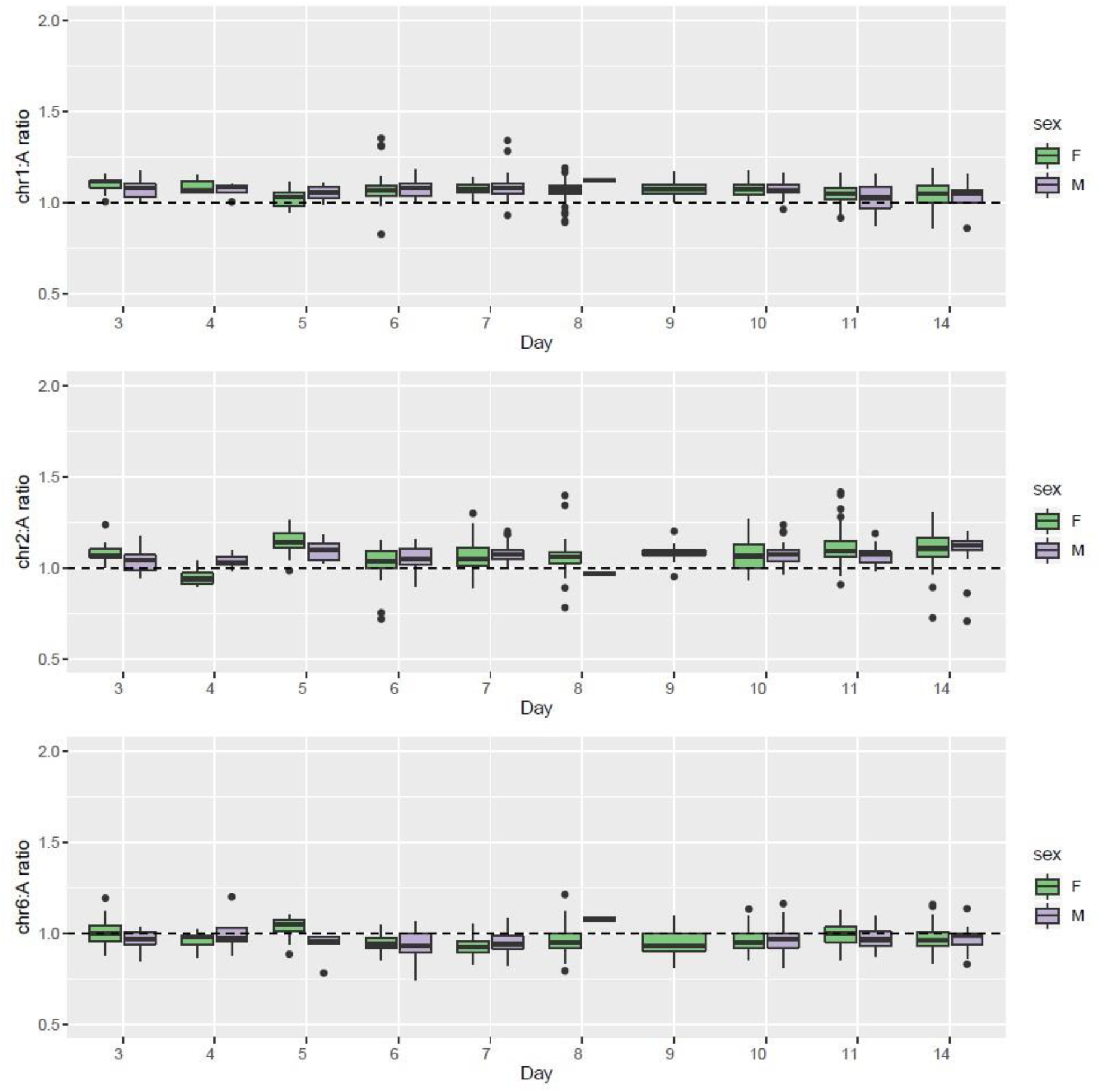
X:A ratio in autosomes. Boxplot showing the distribution of autosome-to-autosome ratios in single-cells across different days of in vitro culture. Shown are representative chromosomes chr1, chr2, and chr6. Center line represents the median, box limits represents upper and lower quartiles, whiskers are 1.5x interquartile range, and points represents outliers.

**Supplementary Figure S3.**
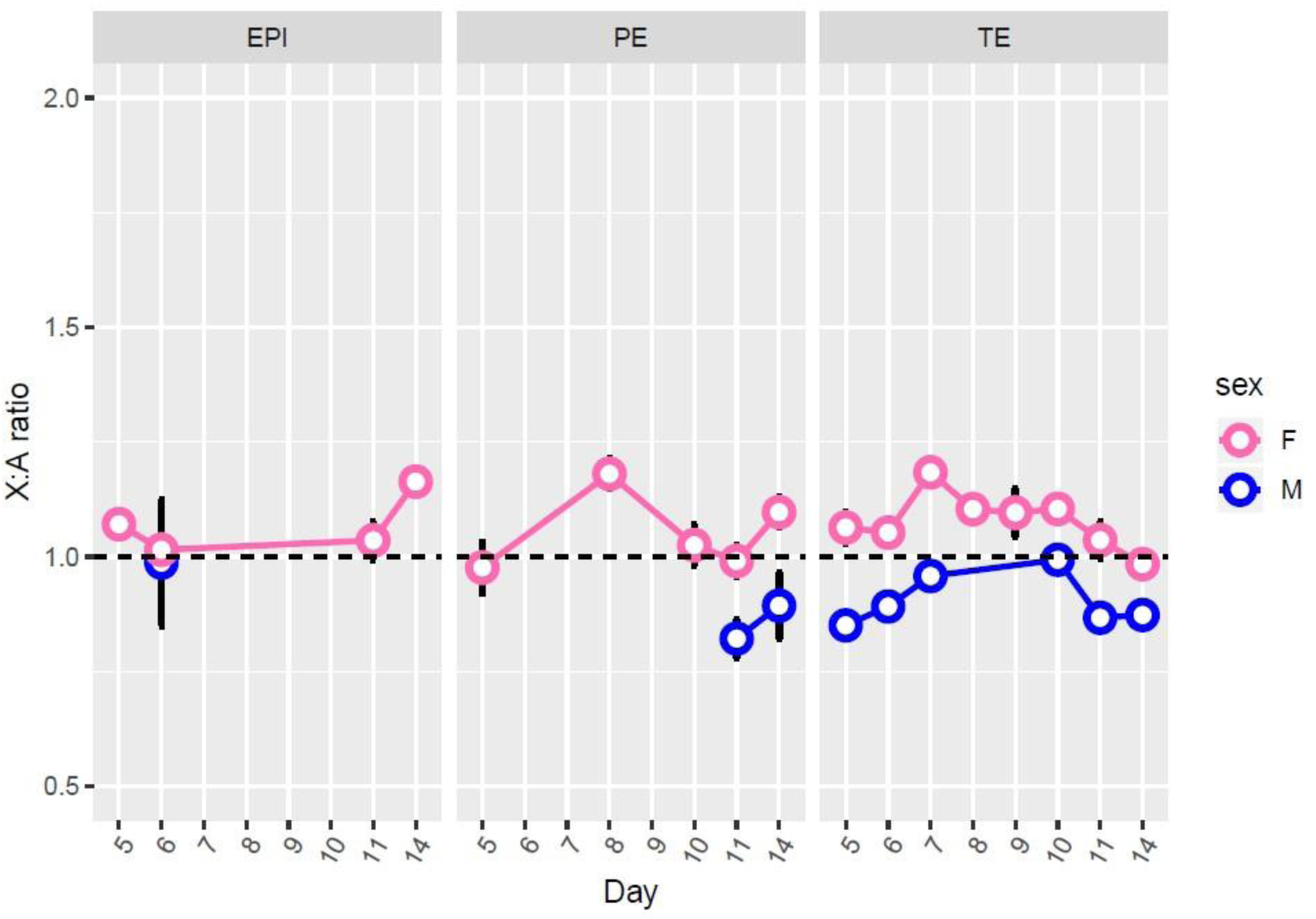
X:A ratio in three lineages. Line plot showing the X:A ratio across separate lineages. Each point represents the median X:A ratio in each study at a particular time point (x-axis) for either female (open circle) and male (open triangle). Error bars represent the standard error. EPI: epiblast; PE: primitive endoderm; TE: trophoblasts.

**Supplementary Figure S4.**
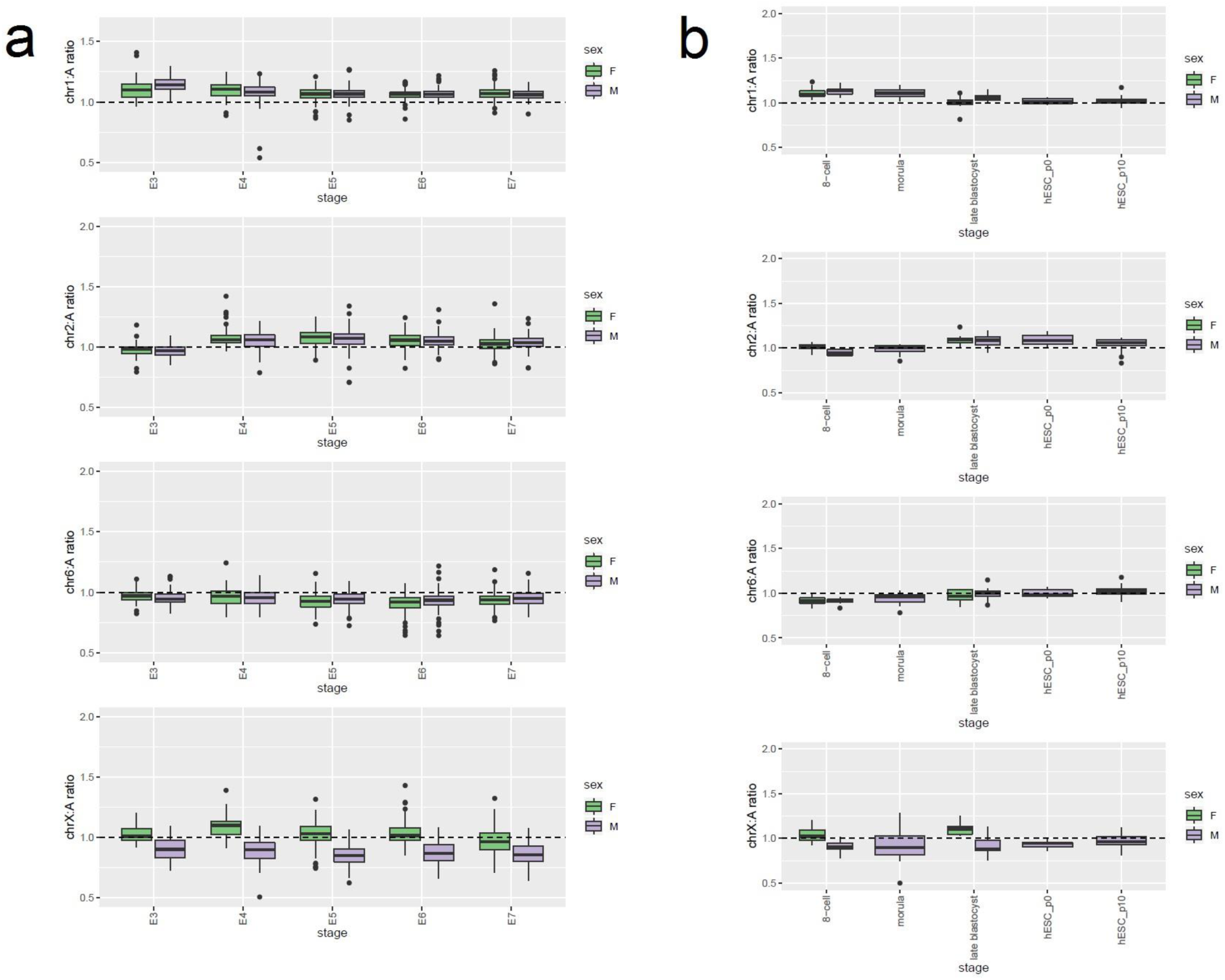
X:A ratio in independent human embryo studies using singlecell RNA-seq. Boxplot showing the distribution of autosome-to-autosome and X-to-autosome ratios in single-cells across different days of in vitro culture in two separate datasets from a) Petropoulos et al. (2016) b) Yan et al (2013). Center line represents the median, box limits represents upper and lower quartiles, whiskers are 1.5x interquartile range, and points represents outliers.

**Supplementary Figure S5.**
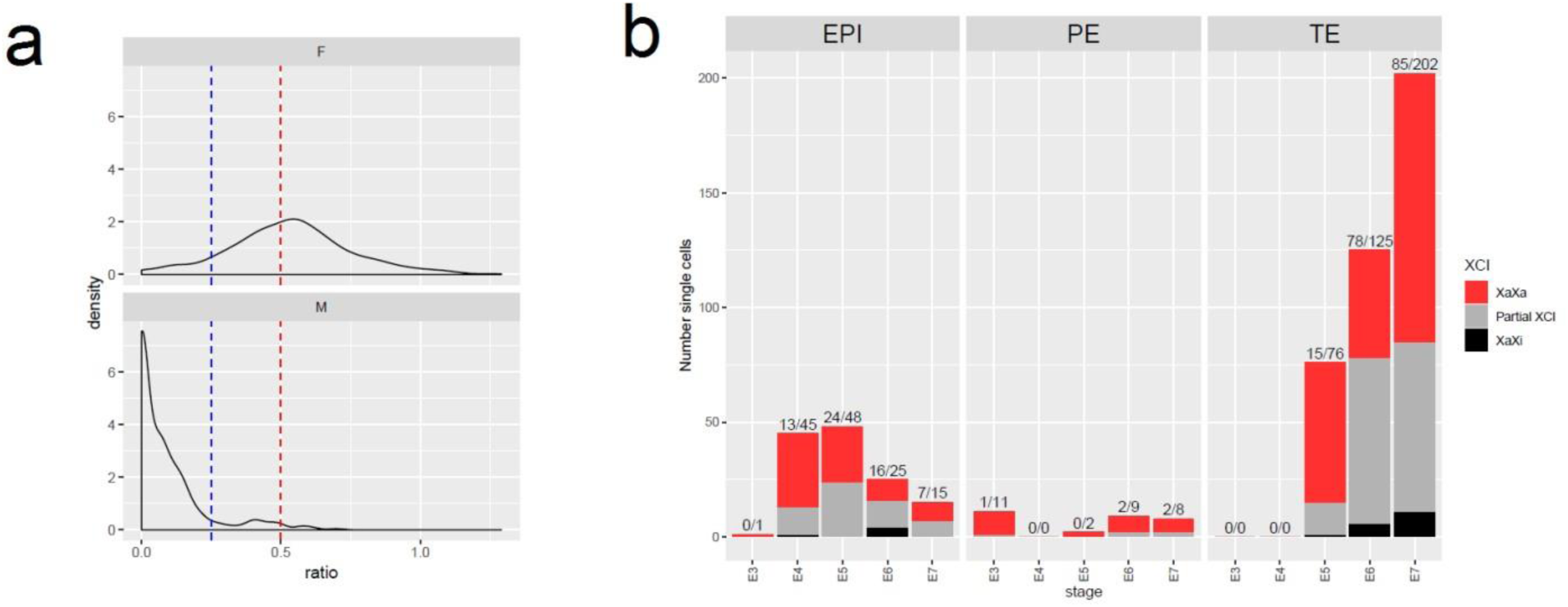
X-inactivation dynamics in Petropoulos et al dataset. a) Density plot showing distribution of biallelic ratio, (the fraction of X-linked biallelic SNPs vs fraction biallelic autosome SNPs). F: female; M: male. Dotted lines show the cutoff for determining monoallelic and biallelic X chromosome status alongside a one-way binomial test (see Methods). X-axis represents the embryonic day, or the number of days in culture. EPI: epiblast; PE: primitive endoderm; TE: trophoblasts. b) Stacked barplot showing the distribution of Xinactivation states across lineages and across developmental days. Numbers above the bars represent the number of partial and complete XCI cells followed by the total number of single cells.

**Supplementary Figure S6.**
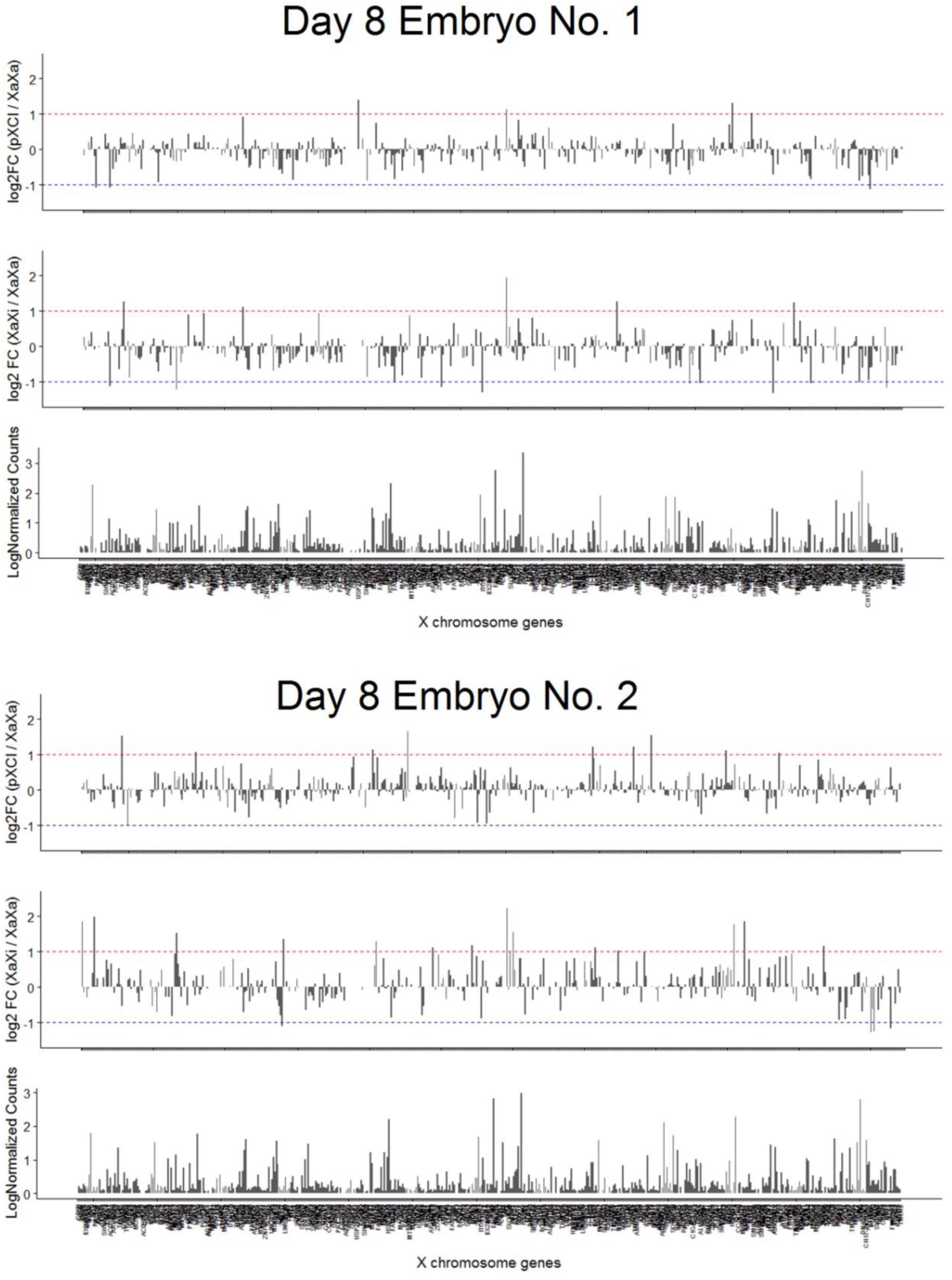
Differential gene expression across the X-chromosome between XaXa, pXCI, and XaXi sister cells. Barplot showing the median log2 fold change of genes between pXCI vs XaXa (top) or XaXi vs XaXa sister cells (middle panel). Bottom panel shows the median log normalized counts for each corresponding gene. This figure is a more comprehensive version from Figure 3b.

